# Wide and Deep Learning for Automatic Cell Type Identification

**DOI:** 10.1101/2020.10.09.328732

**Authors:** Christopher M. Wilson, Brooke L. Fridley, José Conejo-Garcia, Xuefeng Wang, Xiaoqing Yu

**Affiliations:** Department of Biostatistics and Bioinformatics, H. Lee Moffitt Cancer Center & Research Institute, Tampa, FL 33612, USA; Department of Immunology, H. Lee Moffitt Cancer Center & Research Institute, Tampa, FL 33612 Tampa, FL 33612, US

## Abstract

Cell type classification is an important problem in cancer research, especially with the advent of single cell technologies. Correctly identifying cells within the tumor microenvironment can provide oncologists with a snapshot of how a patient’s immune system is reacting to the tumor. Wide deep learning (WDL) is an approach to construct a cell-classification prediction model that can learn patterns within high-dimensional data (deep) and ensure that biologically relevant features (wide) remain in the final model. In this paper, we demonstrate that the use of regularization can prevent overfitting and adding a wide component to a neural network can result in a model with better predictive performance. In particular, we observed that a combination of dropout and *ℓ*_2_ regularization can lead to a validation loss function that does not depend on the number of training iterations and does not experience a significant decrease in prediction accuracy compared to models with *ℓ*_1_, dropout, or no regularization. Additionally, we show WDL can have superior classification accuracy when the training and testing of a model is completed data on that arise from the same cancer type, but from different platforms. More specifically, WDL compared to traditional deep learning models can substantially increase the overall cell type prediction accuracy (41 to 90%) and T-cell sub-types (CD4: 0 to 76%, and CD8: 61 to 96%) when the models were trained using melanoma data obtained from the 10X platform and tested on basal cell carcinoma data obtained using SMART-seq.

## 1 Introduction

Immunology is quickly becoming a popular area of study in cancer research and offers an opportunity to expand our understanding and ability to treat patients. Estimating the immune composition of an individual’s tumor has been the focus of several studies which have developed deconvolution methods [17, 24] to estimate the cellular composition of the tumor micro-environment with bulk RNA expression data. Recently with the advent of single cell sequencing researchers are now able to measure gene expression in individual cells within the tumor-microenviroment and classify cells using heirarchical clustering and correlation-based methods [1, 2, 7]. Cell type classification can be conducted by constructing visualizations such as t-Distributed Stochastic Neighbor Embedding (t-SNE) [28] or Uniform Manifold Approximation and Projection (UMAP) [16] plots to define clusters and assign these clusters to different cell types based on enriched canonical markers. However, a major drawback of this canonical process is that it heavily relies on the researchers’ knowledge on the cell-type-specific signature genes, and it can become arbitrary when making conclusions based on only a handful of genes. Also, the cell type marker genes are cancer type-specific and may not generalize to other datasets [31]. In addition, discriminating between fine immune cell sub-types, such as exhausted CD8 T cells vs. activated CD8 T cells, effector CD4 T cells vs. naive CD4 T cells is a much more challenging task due to the lack of universal marker genes.

Identification of highly specific cell types is now possible with the development of single cell RNA-sequencing technology. However, a challenge in cell annotation in single cell RNA-sequencing is that transcription profiles are difficult to transfer between different platforms. Multiple platforms have been developed for single cell RNA-sequencing including SMART-seq [18], CEL-seq [11], Fluidigm C1 [13], SMART-seq2 [19], and more advanced droplet-based platforms including Drop-seq [15] and 10X Genomics Chromium system [35], etc. The two most commonly used platforms are SMART-seq/SMART-seq2 and 10X. The 10X platform is a droplet-based approach which generates a unique molecular identifier (UMI) at 5’ or 3’ ends to diminish the sequencing reads representation biases due to library amplification. On the other hand, SMART-seq and SMART-seq2 are designed to generate full-length cDNA.

Droplet-based 5’ or 3’-tag methods like 10X can capture much more cells which in turns can give better overview of the heterogeneity within population; while a full-length proposal like SMART-seq is better suited for studies concerned with isoforms, splicing or gene fusion. Due to the differences in how they amplify the mRNA transcripts, the data generated from these platforms are not directly comparable, which presents great challenge to the integrated cell type identification in cross-platform datasets [4, 20, 29, 34]. Therefore, there is a great need for automatic cell identification method that be used across studies, single cell platforms, and cancer types.

While there are many different single cell RNA-sequencing platforms whose results are on different scales and not directly comparable, the underlying gene to gene relationships should be consistent and navigating these relationships may allow for borrowing of information from different technologies. Deep learning brings us the possibility to explore and summarize complex highly non-linear relationships into high-level features from high throughput data sources. Deep learning is a powerful machine learning technique that is often used in visual recognition [12, 14], natural language processing [21, 32], and starting to infiltrate the realm of cancer research [3, 5, 25]. Deep learning learns patterns in data by using neural networks with many layers of nodes which transform the output model of the nodes from the previous layer with non-linear functions. The coefficients output from each node are augmented using gradient descent in order to optimize the prediction error of the network.

Wide and deep learning (WDL) combines a deep neural network with a generalized linear model (GLM) based on a small set of features. Deep learning tends to generalize patterns in the data, while in contrast GLMs may only memorize the patterns in the data. WDL has been shown to be an effective tool in recommender systems [6]. Specifically, we propose utilizing a deep learning model which can leverage large dimensional data (deep), as well as, incorporate a few known biologically relevant genes in the last hidden layer of a neural network to emphasize their biological importance (wide). The wide part of the model allows us make cell type classification more precise and fine tuned to classify more specific immune cell sub-types such as distinguishing activated from exhausted CD8 T cells.

This paper serves two purposes. First it provides some background information about deep learning specifically focusing on regularization methods to avoid overfitting the model. Models are trained, validated, and subsequently used to classify cells from the same dataset. This scenario is realistic since it is possible that some hospitals may not have the resources needed for generating large amounts of data to build their own model. In addition, in many clinical studies the patients’ tumor samples are collected over a fairly long period of time (years) in several batches. Waiting till sample collection is finished before single cell RNA-sequencing data analysis is not realistic. It will be extremely helpful to train a deep learning model using samples collected at earlier time points and subsequently apply it to later samples. Second, this paper provides an illustration of how incorporating known biologically relevant biomarkers can be used to transfer knowledge. In this scenario, we explore the possibility to transfer cell type annotations across different single cell RNA-sequencing platforms, which can help make full use of the enormous publicly available single cell RNA-seq data that are generated by different technologies.

In the methods section, we will describe the data single cell RNA-seq datasets used in study, provide background about deep learning, and wide and deep learning. Then in the results section, we will present results from training and testing the models in the two scenarios. Finally, we make some concluding comments and discussion in the discussion section.

## 2 Methods

### 2.1 Chang Data

Chang et al. [31] conducted droplet-based 10X 5’ single-cell RNA-sequencing on 79,046 cells from primary tumors of 11 patients with advanced basal cell carcinoma before and after anti-PD-1 treatment. In total, RNA profiles from 53,030 malignant, immune and stromal cells, and 33,106 T cells were obtained from single cell RNA-sequencing. The cell types of interest were T cells, B cells, nature killer (NK) cells, macrophages, cancer associated fibroblasts (CAFs), endothelial cells, plasma cells, melanocytes, and tumor cells. The T cells were further classified into regulatory (Tregs), follicular helpers (T_FH_), T helper 17 (T_h_17), naive T cells, activated CD8, exhausted CD8, effector CD8, and memory CD8 T cells (Supplementary Figure 1).

### 2.2 Tirosh data

Tirosh et al. [26] applied SMART-seq to 4645 single cells isolated from 19 freshly procured human melanoma tumors, profiling T cells, B cells, NK cells, macrophages, endothelial cells, CAFs, and melanoma cells. To further analyze the T cell sub-types, we downloaded the log-transformed TPM (Transcripts per Million reads) gene expression values provided by the study and imported them to Seurat [23]. S and G2/M cell cycle phase scores were assigned to cells based on previously defined gene sets [27]using CellCycleScoring function. Scaled z-scores for each gene were calculated using ScaleData function by regressing the S and G2/M phases scores. Shared nearest neighbor (SNN) based clustering method was used to identify clusters based on the first 30 principle components computed from scaled data with resolution = 1. UMAPs were generated using the same principle components with perplexity = 30 and used for all visualization. Clusters were annotated by identifying differentially expressed marker genes for each cluster and comparing to known cell type markers and markers reported by Tirosh et al. From this analysis, we confirmed the cell annotation provided by Tirosh et al., and were able to further identify CD4+ T cells and CD8+ T cells.

### 2.3 Background for Classification Problems with Deep Learning

Deep neural networks (DNN) are able to learn and condense highly non-linear features (genes) into a high level summary through the use of composition of functions. These functions are dot products that undergo a non-linear transformation and are then passed into another function in the next layer. There are three types of layers in a DNN which are input, hidden, and output layers. Each node in the input layer corresponds to the expression of a single gene. Information from the input layer is passed to each node in the a hidden layer which optimizes the weights in a dot product to maximize the cell type classification accuracy. The nodes in the output layer produce the probability that a cell is classified as a specific type. The architecture of a generic DNN with two hidden layers is seen in Figure 1. For multi-class classification the objective function to be minimized is so called cross-entropy function, *H* (*P, Q*),

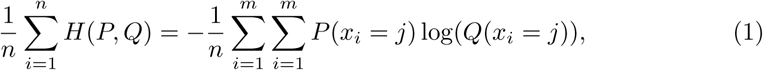

where *m* is the number of cell types, *n* is the sample size, *P* is the target probability distribution and *Q* is the predicted cell type probability distribution. This function is minimized by gradient descent which is computed by iterating the chain rule over all layers of the model. The architecture of a DNN is complex and requires careful tuning. Some examples of tuning parameters in a DNN are number of samples used for stochastic gradient, iterations to train the model, and nodes in each layer [9, 10].

**Figure 1.**
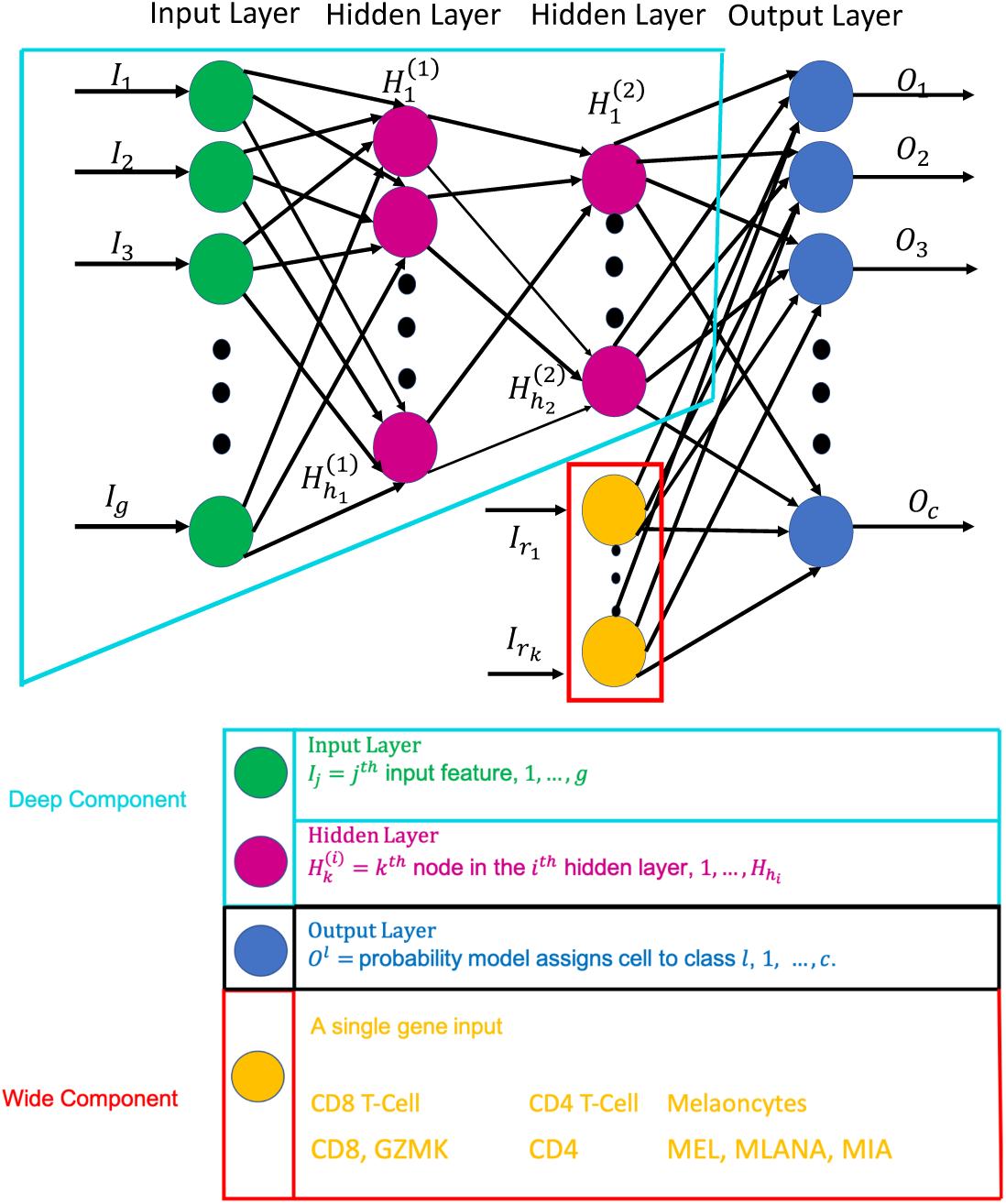
Depiction of a generic wide and deep learning neural network where the wide component is surrounded by the red lines and the deep component is encompassed by the turquoise lines. The deep component equivalent a traditional deep learning model described in sections 2.2 and 3.1. The gene names in yellow correspond to the genes that are used in the wide part of the WDL model in section 3.2.

A challenge to training a deep learning model is to ensure that the results can be generalized to new data sets. One of the simplest ways to prevent overfitting is use to reduce the number of hidden layers or nodes which in turn decreases the number of parameters estimated by the model. Another technique is dropout which randomly deletes a specified proportion of nodes from each layer in the neural network. By deleting different sets nodes in each iteration the model is trained on different sub-networks and becomes less sensitive to the specific weights of nodes. Dropout can speed up the training of a DNN, however, it may require more iterations to train the network. It is recommended that the percentage of nodes to delete from each layer should be between 20-50% [22] Lastly, constraints can be imposed on the weight vector of each node requiring the norm be small. The regularizers work in a similar way to lasso or ridge penalization in a regression setting where there is an additional parameter which changes the influence of the penalization term. The aim is either to keep the value of the weights small or push as many as possible to zero (lasso). Elastic net penalization has also been used which allows for a balance between the *ℓ*_1_ (lasso) and *ℓ*_2_ (ridge) penalty [9, 30, 33].

Understanding which genes were influential to a successful cell type classification model is important for validating the results and can lead to detection of novel genes for future research. Despite many machine learning techniques being seen as ‘black boxes’, there have been efforts to interpret the results. One simple approach to evaluate the importance of a feature is to calculate dot product of consecutive nodes [8]. This was originally proposed for neural networks with a single hidden layer, but we extend this work to a neural work with two hidden layers

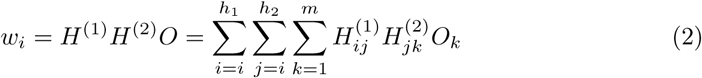

where 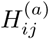 is the coefficient passed from the *i*^*th*^ node of the (*a −* 1)^*st*^ hidden layer to the *j*^*th*^ node in the *a*^*th*^ hidden layer, and *O*_*k*_ probability that a cell is from the *k*^*th*^ cell type. By ranking these weights of each gene we can gain some notion of variable importance in the final classification.

### 2.4 Emphasizing Important Genes for Improved Cell Type Classifiers

Wide and deep learning (WDL) involves merging a set of features, wide component, with the last hidden layer in a DNN, deep component. Adding these features in the final step will ensure that they are emphasized in the model, since they may be lost due to dropout or assigned with small weights. The wide component is a generalized linear model where the input is a set original features. Wide components tend to memorize the patterns the data, while deep components can generalize non-linear patterns. The architecture of a WDL model is shown in Figure 1. In this study, specific genes that are exclusively expressed by a particular cell type are added to the last hidden layer forcing the model to emphasize them more. This may allow a DNN to produce a more accurate classify cells model than a model constructed with only a deep part, especially in scenarios where the data are obtained from different platforms or cancer types.

## 3 Results

### 3.1 Neural Network Tuning

In this section, we want to describe how the hyperparameters (number of nodes, regularization, dropout) were selected. Traditional deep learning models with two hidden layers were constructed with no regularization (No Dropout), 20% dropout for both hidden layers (Dropout Only), 20% dropout and an *ℓ*_1_ regularizer for both hidden layers (Dropout + *ℓ*_1_), and lastly 20% dropout and an *ℓ*_2_ regularizer for both hidden layers (Dropout + *ℓ*_2_). A grid search was employed where the first hidden layer could have 1, 5, 10, 25, 50, 75, 100, 500, 1000 nodes and the second hidden layer could have 1, 5, 10, 25, 50, 75, 100, 500 nodes.

Figure 2 shows the training and validation accuracy (left) and loss (right) when there was the same architecture for each model, and a numerical summary is provided in Table 1 for the architecture leading to the best validation accuracy. Notice that with No Dropout, Dropout Only, and Dropout + *ℓ*_1_ the validation loss increases as the model is trained. On the other hand, the validation accuracy and validation loss remains consistent with the training loss as epochs increase for Dropout + *ℓ*_2_, suggesting that even if the model is trained with an excessive number of iterations that the model performance will not suffer heavily from overtraining. While the No Dropout and Dropout only models had the lowest validation loss and nearly 100% training accuracy, they are undoubtedly overtrained and will likely not generalize well for future data. Based on these finding, all models discussed in the remainder of this paper will be constructed with dropout and *ℓ*_2_ regularization especially since there is not a significant difference between the testing accuracies of the four methods.

**Table 1.** Summary of the parameter selection, validation, and testing accuracy for the naive, and wide and deep learning models. The architecture was selected based on the model that highest accuracy when classifying cells in the validation data set. The testing accuracies arise from training a model with the specified architecture with training + validation datasets and testing on previously unused test set.

|  | Validation Accuracy | First Layer | Second Layer | Testing Accuracy |
| --- | --- | --- | --- | --- |
| No Dropout | 94.4 | 100 | 50 | 95.2 |
| Dropout Only | 94.2 | 100 | 75 | 94.5 |
| Dropout + $\ell_2$ | 94.3 | 100 | 75 | 93.8 |
| Dropout + $\ell_1$ | 92.9 | 75 | 100 | 92.2 |

**Figure 2.**
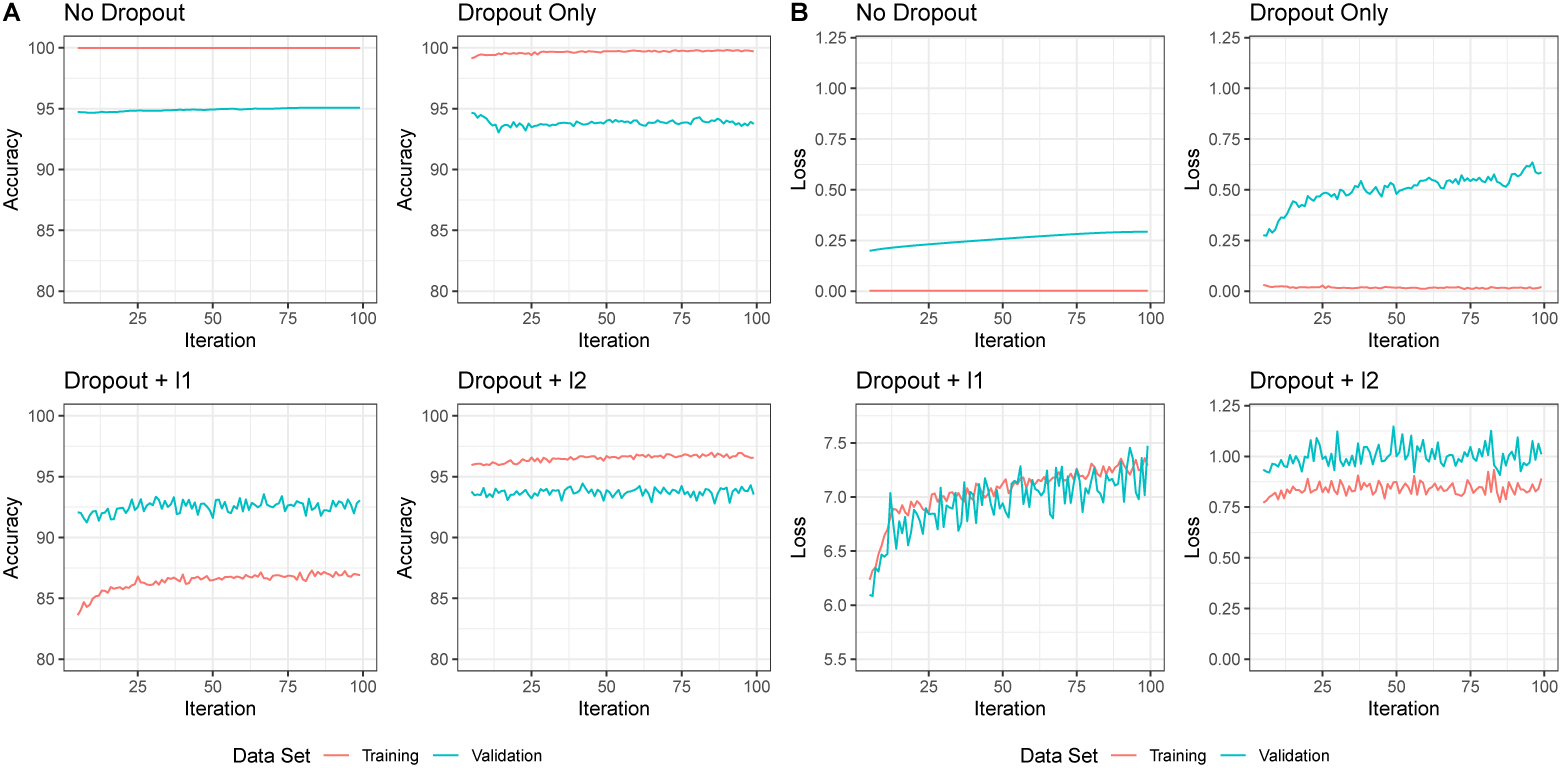
Comparison of the accuracy (A) and categorical cross-entropy loss (B) of four models from section 3.1 with varying methods of regularization and drop out. The plots are for iteration number 5 through 100. The red and turquoise lines correspond to the performance on the training validation set, respectively.

The overall accuracy of the Dropout + *ℓ*_2_ model is 93.8%, with the prediction accuracy of individual cell types ranging from 83 to 100% (Figure 3A). T-cell sub-types are similar in gene expression profiles and are difficult to distinguish. T-cell sub-type classification is commonly done as a second stage of classification where only the T-cells are considered [31]. Figure 3A shows that using a deep learning framework, each T-cell sub-type is classified with at least 83% accuracy, and 5 out of the 7 T-cell sub-types had greater than 91% percent accuracy, and the misclassified cells were classified as another type of T-cell. In single cell RNA-sequencing, the separation between activated CD8 and exhausted CD8 T cells are particularly difficult. The exhausted cells are considered as chronically activated and they also highly express the activation markers such as TNF and IFNG. The subtle difference between activated and exhausted CD8 T cells is the overexpression of exhaustion markers such as TIGIT and HAVCR2. Our deep learning model with Dropout+ *ℓ*_2_ setting was able to capture these genes and ranked their importance as 4^*th*^ and 75^*th*^ in total of 22,890 genes (Figure 3B, Supplementary Table 1). In addition, the model also put high emphasis on the genes that are typically over-expressed in tissue-resident memory cells, such as LAYN and CXCL13 (Figure 3A), which is consistent with Chang et al. original findings. The cell type classification accuracies for No Dropout, Dropout only, and Dropout + *ℓ*_1_ are included in Supplementary Figure 2.

**Figure 3.**
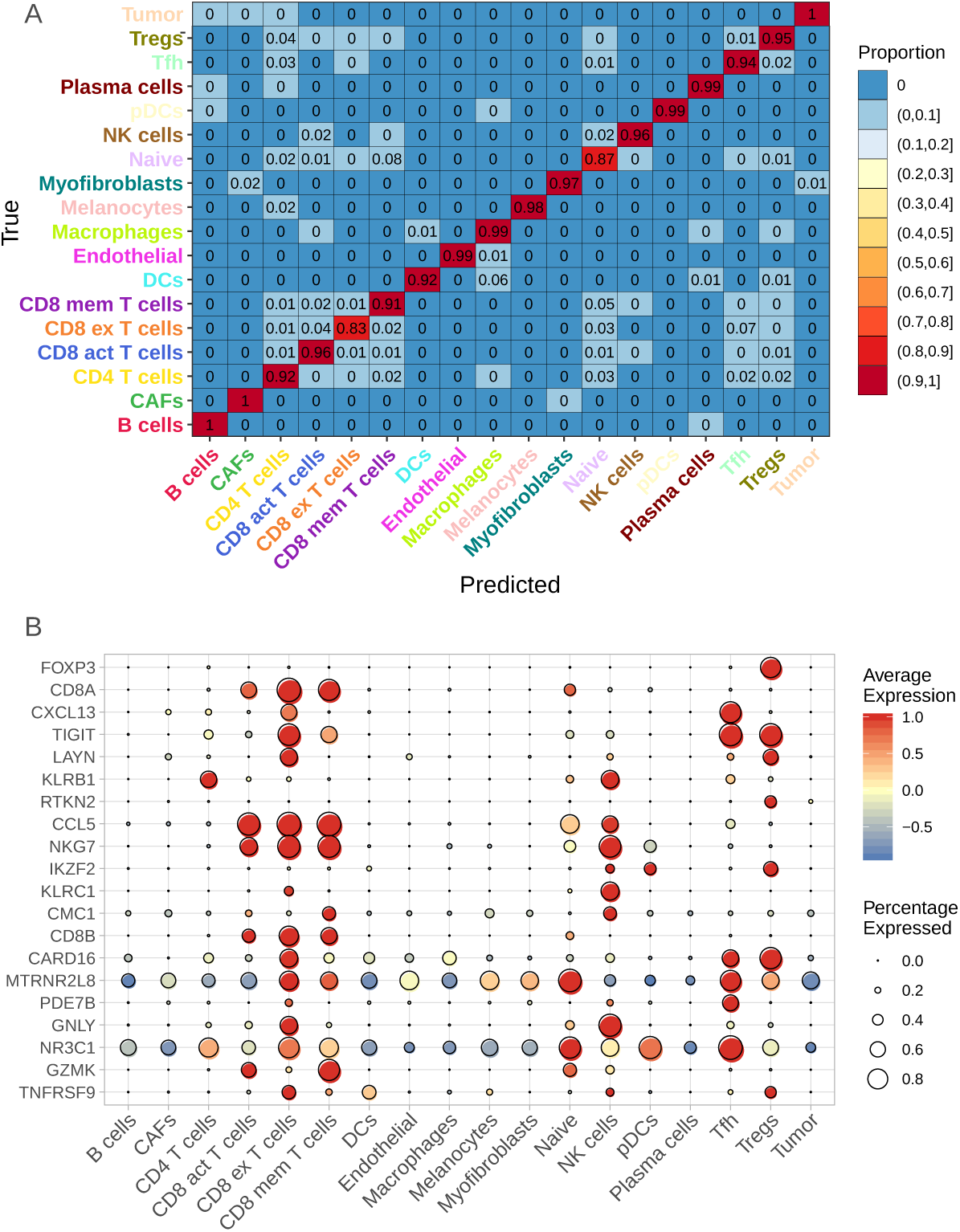
Heatmap of accuracies by cell type for a deep learning model (A), trained and tested on the Chang dataset, with two hidden layers with 100 and 75 nodes respectively, and 20% dropout and *ℓ*_2_ regularization. Average expression of the top 20 most influential genes by cell type (B) where the size of the dot corresponds to the proportion of cells that express this gene and color ranging from blue to red indicating low to high average expression.

### 3.2 Testing on Different Datasets

Both naive and WDL learning models were constructed using the melanoma data generated by Chang for training and basal cell carcinoma data produced by Tirosh for testing the models. Both models were constructed with 100 and 75 nodes for the first and second layers respectively. A comparison of the true cells types and the predicted cell types from the WDL model are shown in Figure 4A and 4B. The naive model had an overall accuracy of 41% which is partly due to the model not classifying any cells as CD4 T cells (Figure 4C). Another discrepancy is that a majority of melanoma cells were classified as CAFs (62%), but the silver lining is that the naive model can distinguish tumoral from stromal cells. A large percentage of NK cells were also classified as CD8-T cells (45%) which is not surprising based on the similarity in cellular function and location in the UMAP in Supplementary Figure 1.

**Figure 4.**
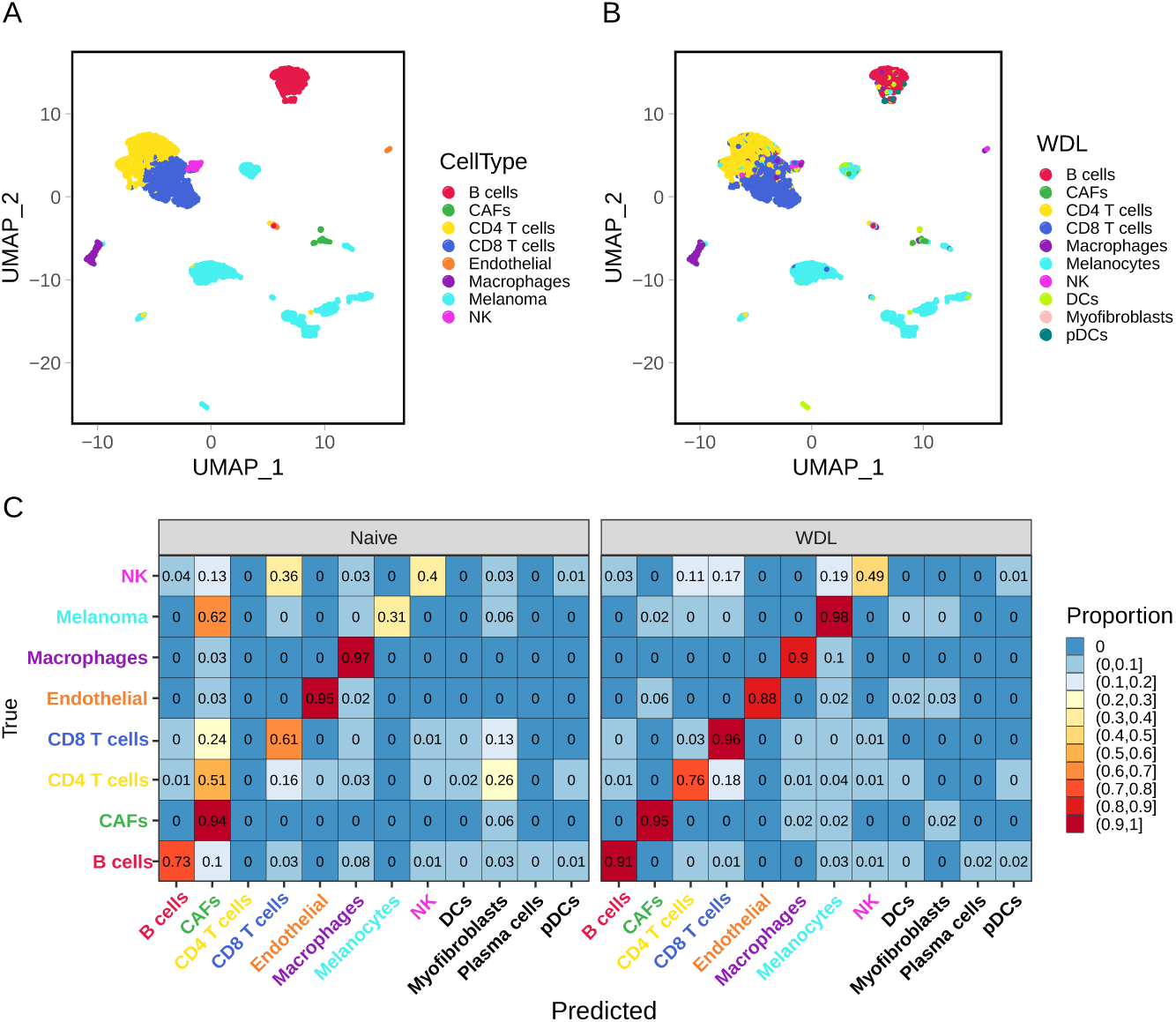
Comparison of the true cell types (A) and predicted cell types from a WDL model (B). Side-by-side comparison of the accuracy by cell type for the naive and WDL models (C).

With the addition of the 8 markers listed in Figure 1, the WDL model can better discriminate sub-types of T cells, CD8 T cells and CD4 T cells, with an accuracy of 95.8 and 75.6% respectively (Figure 4B right) and obtained an overall accuracy of 90.1% accuracy (Supplementary Figure 3). The classification accuracy by cell type ranges from 48.6% for NK Cells to 95.8% for CD8 T-cells with 6 of the 8 cell types having greater than 87% accuracy (Figure 4B right). A majority of the misclassified CD8 T-cells were classified classified within the same major cell type. Classification of melanoma cells saw the largest increase in accuracy from 31% to 98%.

In order to understand why the models perform differently, we focus on the differences between specific markers that were highly influential in each model. The importance of the top 20 markers and their importance are displayed in the average expression profiles for the each cell type in Figures 5A and 5B. A total of 8 markers were included in the wide part of the WDL model, and these have by far the largest importance in the model, and four of these (CD8A, CD8B, GZMK, and IL7R) which are all important for identifying CD8 or CD4 T-cell. Thus the importance of these markers explains why the model was able to classify CD8-T cells with an accuracy of 61%. Five out of the top 12 most important markers from the deep component of the WDL model were in the top 20 markers in the naive model with slightly larger weights in the WDL model. Additionally, there were no melanoma markers in the top 20 most influential genes in the naive model leading to many of the melanoma cells being labels as CAFs. Figures 5, Supplementary Figure 4 and Supplementary Table 2 show that are several genes that are highly influential, yet are not expressed in many cells in the training or test set, such as EOMES, SH2D1B, ENC1, VCAM1. This further illustrates a challenge understanding the importance of markers only a small subset of cells in a particular cell type may express these marker yet the model found them to be influential. Chang identified, after restricting data to only T-cells, EOMES as a marker to distinguish from CD4 memory T-cells other T-cell sub-types, while both deep learning models were able to identify this as in important marker without subsetting the data by major cell type (Supplementary Figure 4).

**Figure 5.**
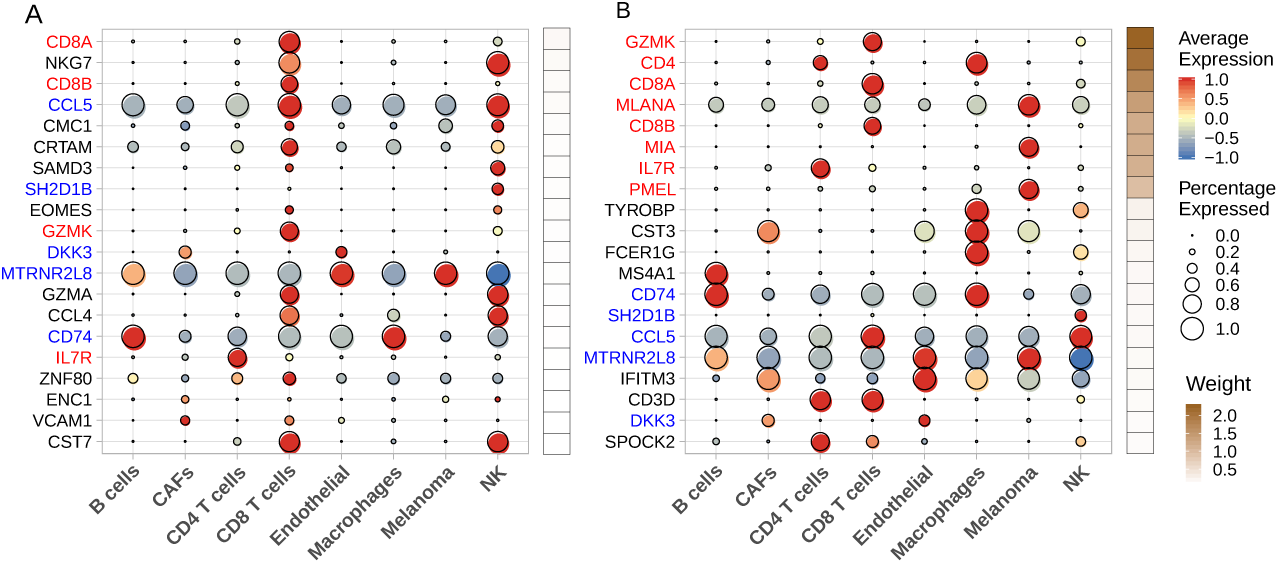
Dot plots for the Tirosh data using both naive (A) and WDL (B) with gene importance weights increasing with brown color scale. The genes names highlighted in red correspond to genes that were included in the wide part of the WDL model, and the blue gene names correspond to the genes that were not in the wide part yet were influential in both the naive and WDL models. Full list of genes and weights is included in Supplemental Table 2.

## 4 Discussion

WDL presents an opportunity to use a small set commonly known biological markers for cell type classification to allow models to be slightly less data driven. We have illustrated a substantial increase in overall accuracy (41 to 90%) and for T-cell sub-types (CD4 increased from 0 to 76% and CD8 increased from 61 to 96%). We have demonstrated that this can allow for training and testing of models from data obtained from different platforms and types of skin cancer, and even when the target classifications are not the same. Further refinement for classification of fine T-cell sub-types is needed to address questions such as ‘how strong is the CD8 T-cell response to a tumor?’, i.e. determine the proportion of CD8 T-cells that are exhausted, which are very relevant in cancer research. Additionally, there is a great need to develop systems to transfer knowledge across cancer types. WDL allows the opportunity to address this by including general set of genes for cell type classification and avoiding adding data/cancer specific markers as shown in section 3.2.

In addition to adding a wide component to a DNN there is need for careful consideration for how model is trained to avoid the memorization of data. While dropout is a great tool for making deep learning models more generalizble there are many applications where there is a need for additional steps to avoid overfitting. Regularization is computationally intensive, but makes deep learning models for generalizable to test datasets. Models can very easily be overtrained but a combination of dropout and *ℓ*_2_ regularization can provide a loss function that is stable across the training iterations. Another challenge for deep learning in general is the randomness in the initialization of node weights, dropout, and batches can lead to dramatically different performances for models that are tuned in the same manner and data. Studying an ensemble of deep neural networks could help study the stability of the models and comparing the most important biomarkers in each model can provide further confidence that the markers that are highly influential. Identifying these genes can help clinicians understand commonality between immune cells behavior across cancer types providing better insight and treatment of the cancers themselves.

## Declarations

The authors declare that they have no known competing financial interests or personal relationships that could have appeared to influence the work reported in this paper.

## Authors’ contributions

All authors read and approved the final manuscript. CW, BLF, JRC, XW, and XY conceived the study. CW, XW and XY designed the algorithm, performed the analyses, interpreted the results and wrote the manuscript.

## Acknowledgments

The authors would like to thank Colleagues at Department of Biostatistics and Bioinformatics at Moffitt Cancer Center for providing feedback.

