## Supplementary material for "Wide and Deep Learning for Automatic Cell Type Identification": Supplementary.Figures.docx

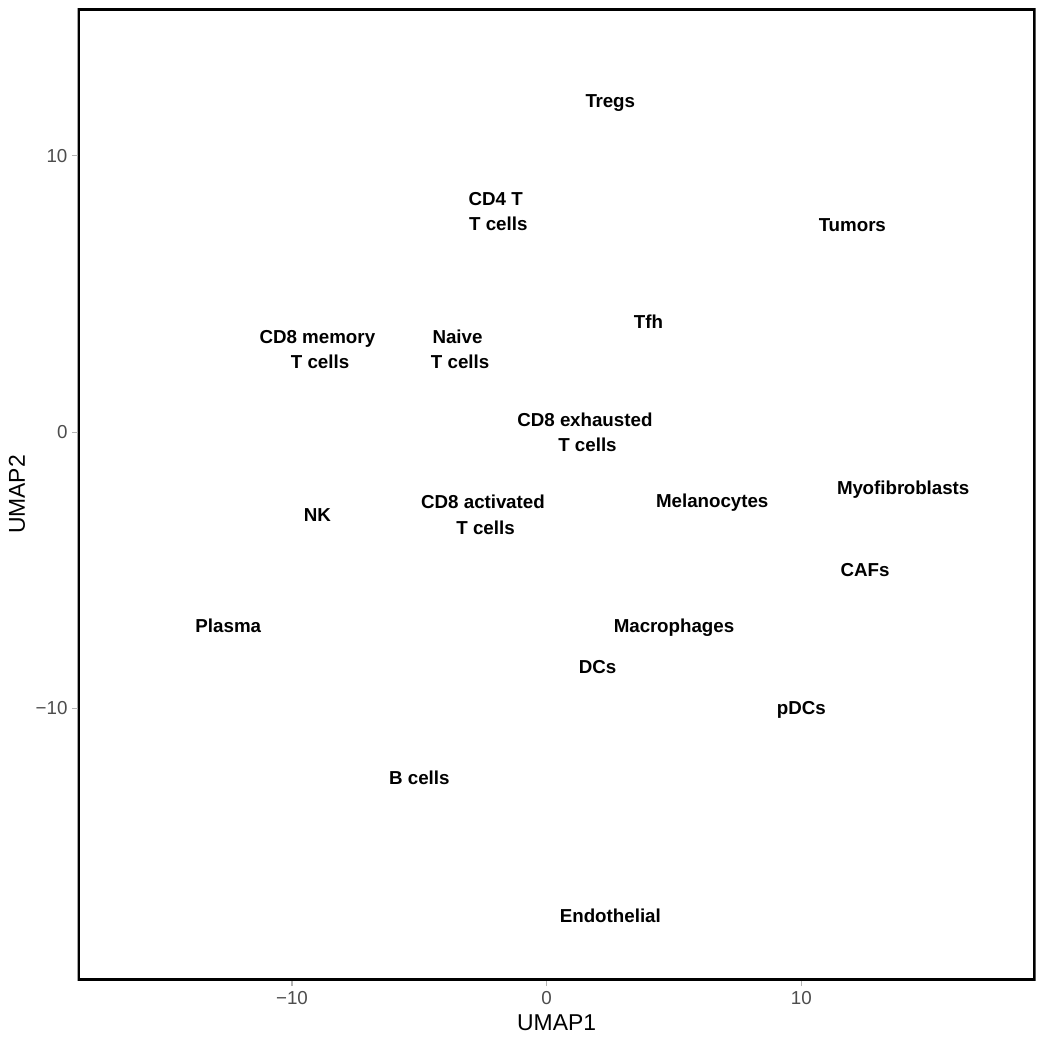

Supplementary Figure 1. UMAP of the Chang dataset colored by cell type.

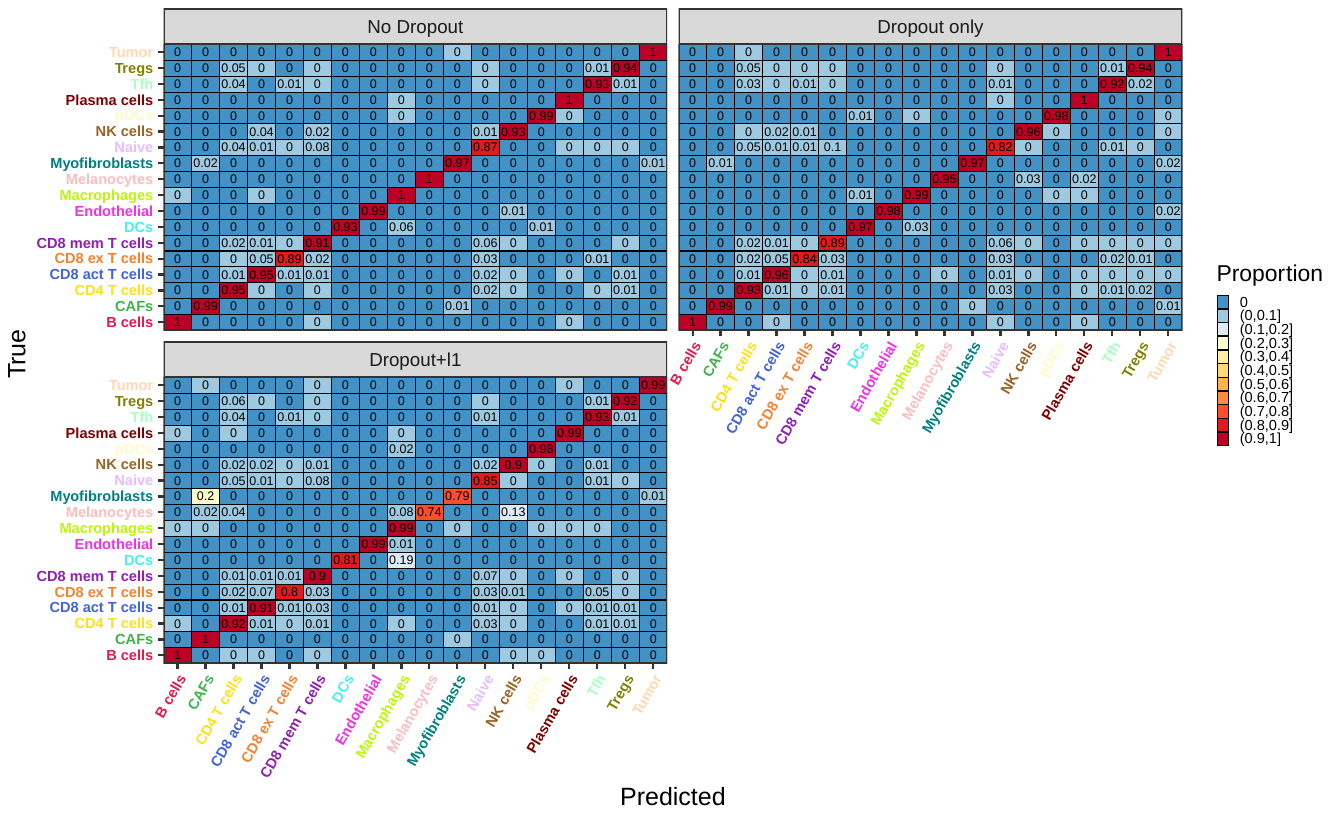
Supplementary Figure 2. Heatmaps of the performance by cell type of models trained with no dropout (No Dropout), 20% dropout (Dropout Only), and 20% dropout and $l1$regularization (Dropout + $l1$).

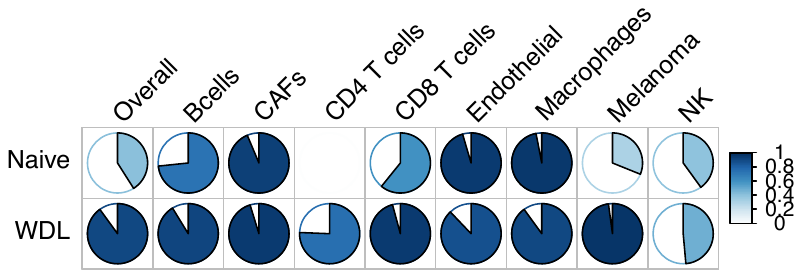

Supplementary Figure 3: Pie chart representations of the overall and by cell type accuracies for both naive and WDL model.

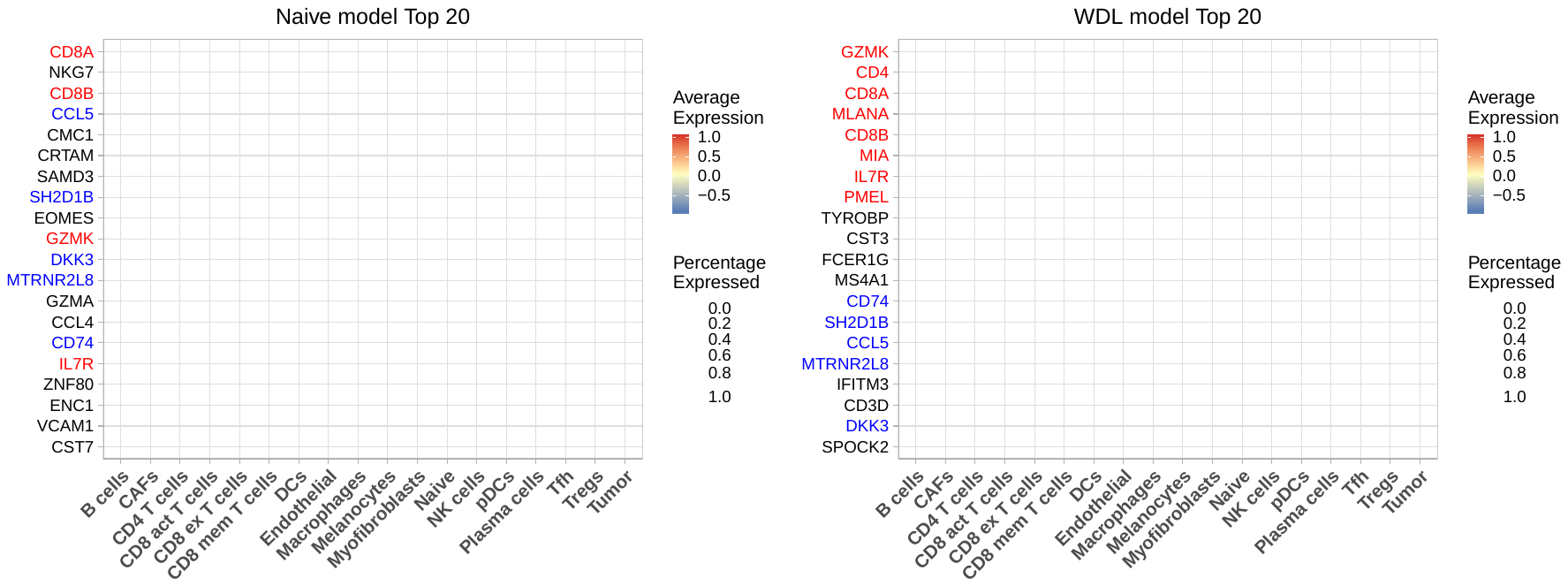

Supplementary Figure 4: Dot plots for the Chang data using both the naive (A) and WDL (B) with gene importance weights increasing with brown color scale. The genes names highlighted in red correspond to genes that were included in the wide part of the WDL model, and the blue gene names correspond to the genes that were not in the wide part yet were influential in both the naive and WDL models. Full set of genes and weights are included in Supplementary Table 1.

**Supplemental Tables**

Supplemental Table 1: Influence of each gene for Section 3.1 for classifying 17 cell types, top 20 genes were included in Figure 3B.

Supplemental Table 2: Influence of each gene for Section 3.2 for classifying 8 cell types, top 20 genes were included in Figure 5. The 8 genes included in the wide component are highlighted in red.
